# Sensory responses in dorsolateral striatum are modulated by motor activity in a dopamine-dependent manner

**DOI:** 10.1101/2022.05.03.490413

**Authors:** Roberto de la Torre-Martinez, Maya Ketzef, Gilad Silberberg

**Affiliations:** Department of Neuroscience, Karolinska Institutet, Stockholm 17177, Sweden

**Keywords:** Striatum, *in vivo*, whole-cell recordings, awake, optogenetics, sensorimotor, medium spiny neurons, whiskers, Parkinson’s disease, basal ganglia, active sensing.

## Abstract

The dorsolateral striatum (DLS) receives excitatory inputs from both sensory and motor cortical regions and is involved in sensory and motor functions. In cortical regions, sensory responses are altered by motor activity, however, it is not known if such sensorimotor interactions also occur in the striatum and how they are modulated by dopamine (DA). To determine the impact of motor activity on striatal sensory processing, we performed *in vivo* whole-cell recordings in the DLS of awake mice during the presentation of tactile stimuli. Striatal medium spiny neurons (MSNs) were activated by both whisker stimulation and spontaneous whisking, however, responses to whisker deflection during ongoing whisking were attenuated. DA depletion reduced the representation of whisking in direct-pathway MSNs, but not in those of the indirect-pathway. Furthermore, DA depletion impaired the discrimination between ipsi- and contralateral sensory stimulation in both direct- and indirect-pathway MSNs. Our results show that sensory responses in basal ganglia circuits are modulated by motor activity and that both processes are dopamine- and cell type-dependent.

## Introduction

Sensory processing in various cortical regions has been shown to be modulated by motor activity ^1–3^. While locomotion reduces the magnitude of auditory responses in auditory cortex ^4–6^, visual responses in visual cortex are enhanced ^7–9^. In the somatosensory cortex, tactile responses to whisker deflection are attenuated during active whisking ^1,10,11^. These studies show that sensory and motor processes are tightly linked at the cortical level, yet it is not known how such sensorimotor interactions are represented in downstream regions. The whisker system is an essential sensory apparatus in rodents, utilizing an active sensing framework where whisking and tactile sensation interact to represent the location and identity of objects ^12^. The dorsolateral striatum (DLS) receives dense monosynaptic excitatory input from sensory cortical regions, including the barrel field, and is involved in sensory representation ^13–18^. Striatal medium spiny neurons of both the direct- and indirect pathways (dMSNs and iMSNs, respectively) respond to whisker stimulation ^16,19–23^. Importantly, MSNs have distinct representations for contralateral and ipsilateral tactile inputs ^16,23^, a feature that is lost following dopamine (DA) depletion, as shown in anesthetized mice ^23^. The DLS also receives axonal projections from motor cortical regions ^14,19,22,24,25^. Such convergence of inputs from both somatosensory and motor cortices suggests that sensory responses are modulated by motor activity also at the striatal level.

Parkinson’s disease (PD) is a neurodegenerative motor disorder associated with the progressive death of midbrain dopaminergic neurons of the substantia nigra pars compacta, a region that, among other structures, densely innervates the striatum ^26,27^. As a result, PD patients typically present motor deficits such as muscle rigidity, tremor, or bradykinesia ^28–30^, some of which are caused by abnormal activation of dMSNs and iMSNs ^23,31–41^. In addition to these motor impairments, PD patients often exhibit somatosensory symptoms such as deficits in the perception of thermal, pain, proprioceptive, and tactile stimuli, including impairments in bilateral tactile discrimination ^30,42–45^.

Here we examine how tactile sensory integration is modulated by motor activity, and how such sensorimotor interactions are affected in a PD mouse model. To achieve this, *in vivo* whole-cell recordings in awake mice were performed in the DLS while bilateral whisker stimulation was delivered in control (DA intact) and DA depleted mice. Responses to sensory stimulation were recorded during different behavioral states, from identified MSNs. Our results show that whisker-related motor activity was correlated with depolarization of MSNs, however, responses to whisker stimuli were attenuated during whisking. DA depletion affected both motor and sensory responses in a cell type-specific manner. Thus, our study provides novel insights into the network mechanisms underlying striatal sensorimotor processing and its impairment in PD.

## Results

### *In vivo* whole-cell recordings from identified MSNs in awake mice

To study sensory processing in MSNs in behaving mice and how it is affected by motor activity, we performed *in vivo* whole-cell recordings from identified dMSNs and iMSNs in the DLS of awake, head-restrained mice (Figure 1A, and 1C). Cell identification during unguided patch-clamp striatal recordings is challenging since the electrophysiological properties of dMSNs and iMSNs are not distinct enough to allow their classification. To overcome this limitation, we used the “*optopatcher*” ^46^ in transgenic mice expressing Channelrhodopsin (ChR2) in either dMSNs or iMSNs resulting from a cross between reporter mouse line (Ai32) with either D1- Cre ^47,48^ or D2-Cre mouse lines (Figure 1B), respectively. We verified the selective expression of YFP-ChR2 in dMSNs or iMSNs by fluorescent labeling of striatonigral and striatopallidal projections, respectively (Figure 1B and 1C). This approach allowed us to identify dMSNs and iMSNs in real-time during recordings using focal photostimulation through the patch pipette (Figure 1D). MSNs expressing ChR2 responded to blue light pulses with immediate and robust depolarization, usually leading to action potential (AP) discharges, whereas ChR2-negative MSNs did not respond to the photostimulation (Figure 1D). The recorded neurons were filled with biocytin to allow for post-hoc visualization of their location in the DLS and morphological reconstruction (Figure 1C). Whisker movements were simultaneously tracked using an infrared LED sensor, allowing the definition of quiescent (Q) and whisking (W) epochs (Figure 1E).

**Figure 1.**
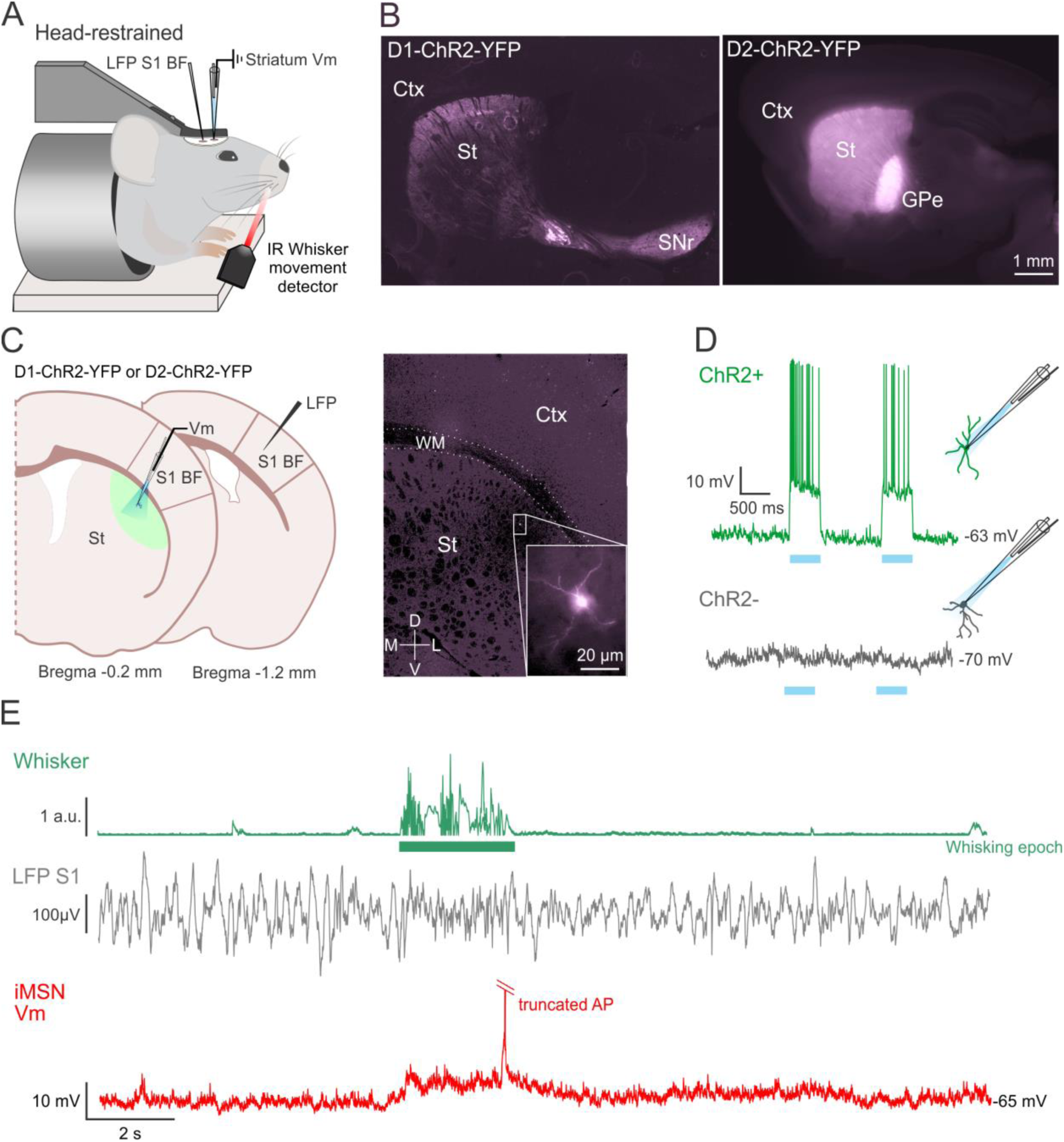
Intracellular recordings from opto-tagged MSNs in the dorsolateral striatum of behaving mice. ((A) Schematic drawing showing the head-restrained mouse. Whole-cell recordings were performed in the DLS with the optopatcher for on-line optogenetic classification parallel to a LFP recordings in S1 BF. Whisking was monitored using a non-contact infrared LED- photodiode. (B) MSNs opto-tagging was obtained using either D1-ChR2-YFP (labeling dMSNs, left) or D2-ChR2-YFP (labeling iMSNs, right) mice. Typical projections from dMSNs in D1-ChR2- YFP mice to SNr and from iMSNs to the GPe in the D2-ChR2-YFP mice are evident. Scale bar, 1 mm. (C) Schematic representation showing intracellular and LFP recording locations from Bregma (left). The image and magnified inset (right) show an example of a biocytin-filled MSN from DLS following *in vivo* whole-cell recording in behaving mice. Inset shows the same cell in higher magnification. Scale bar, 20 μm. (D) Opto-tagging of MSNs using the optopatcher. Image shows depolarizing responses to photostimulation in D1-ChR2-YFP mouse (ChR2+, top) for the duration of the stimulation. Negative cells (ChR2−, bottom) did not respond with depolarization to the light stimulation. (E) Example trace showing spontaneous membrane potential activity in an opto-tagged dMSN from DLS of behaving mouse in control conditions. Membrane potential (Vm, red) of the neuron was recorded simultaneously with measurement of whisker activity (Whisker, green) and LFP activity in BF (LFP S1, black) Abbreviations: Local Field Potential (LFP), Barrel field somatosensory cortex S1BF, infrared light (IR), Substantia Nigra pars reticulata (SNr), Globus Pallidus externa (GPe), Cortex (Ctx); White Matter (WM), Striatum (St), Channelrhodopsin (ChR2).

### DA depletion reduces spontaneous activity in dMSNs

To investigate the effects of DA depletion on the membrane properties of striatal MSNs, we obtained whole-cell recordings from DA-intact (control) and DA-depleted awake mice (Figure 2A, and 2C). DA-depleted mice received a unilateral injection of 6-hydroxydopamine (6- OHDA) in the medial forebrain bundle (MFB) that produced a nearly complete depletion of DA innervation in the striatum of the injected side (Figure 2A, and 2B). We verified the absence of DA by immunofluorescence staining for tyrosine hydroxylase (TH) (Figure 2B).

**Figure 2.**
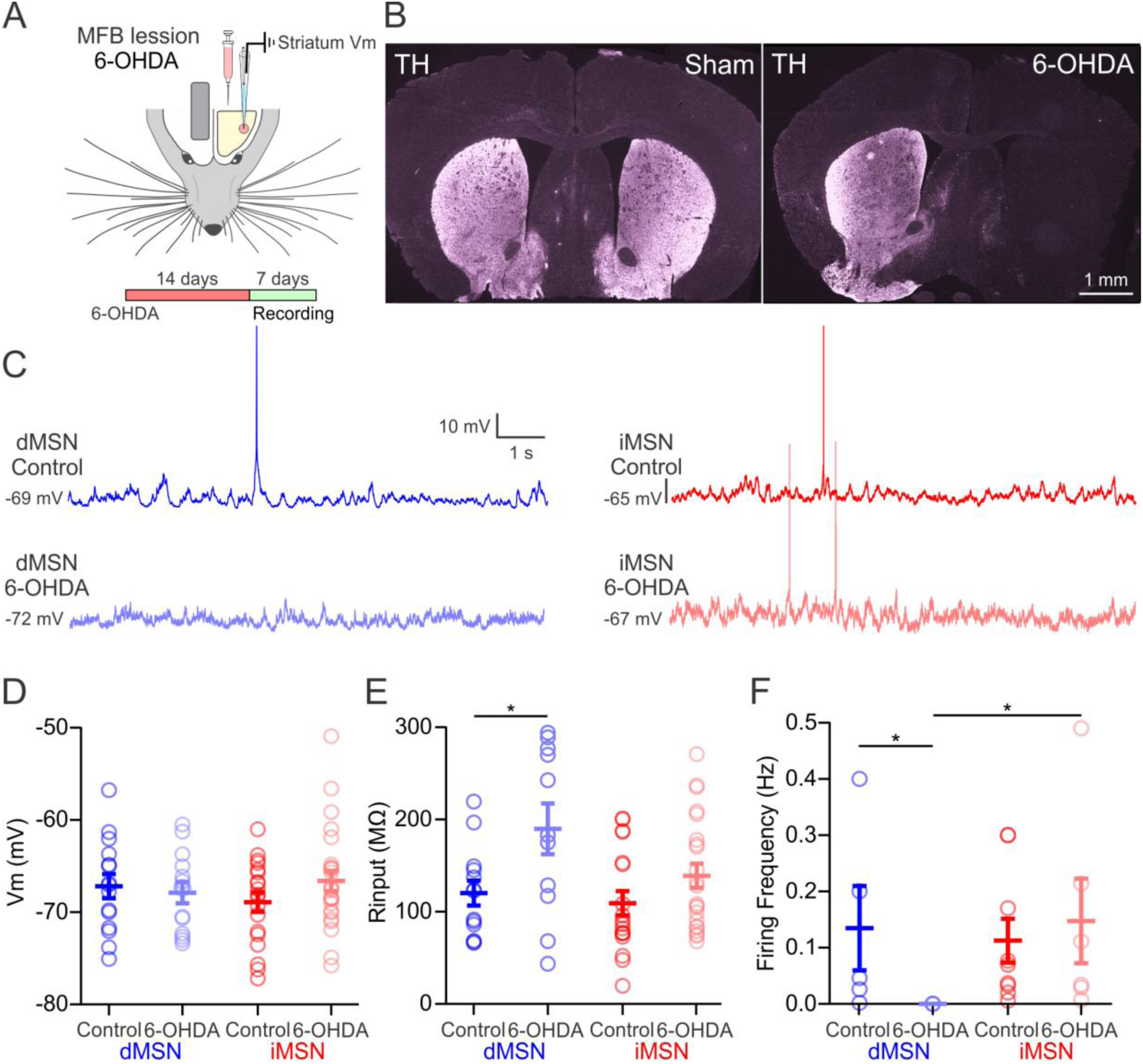
Membrane properties of MSNs in control and 6-OHDA lesioned mice. ((A) Illustration showing the experimental setting, the timeline of 6-OHDA MFB lesion and striatal whole-cell recordings in the same hemisphere at least 14 days following the lesion. (B) Coronal sections showing the striatal hemispheres of sham (left) and 6OHDA-lesioned (right) mice, stained for TH expression. Fluorescence was strongly reduced in the 6- OHDA treated hemisphere compared to the control hemispheres. (C) Example traces showing spontaneous membrane potential activity in dMSNs (blue) and iMSNs (red) in control condition (dark color) and 6-OHDA lesioned (light color) mice. (D) No difference in membrane potential of dMSNs and iMSNs in control and 6-OHDA mice. (E) Input resistance of dMSNs and iMSNs were similar in control conditions. After 6-OHDA lesion, input resistance increased in the dMSNs but not iMSNs. (F) Spontaneous AP discharge frequency of dMSNs and iMSNs showed no differences in control condition. After 6-OHDA lesion, dMSNs were not spiking spontaneously (0/14) while iMSNs did not change their firing frequency. For all panels, data are presented as mean ± SEM. control iMSNs are in dark red, control dMSNs are in dark blue, 6-OHDA lesion dMSNs are in light blue, and 6-OHDA lesion iMSNs are in light red. p < 0.05. Spontaneous firing proportions were analyzed using Fisher’s exact test. * p < 0.05.

The analysis of whisking activity revealed that mice engaged in whisking around 15% of the time throughout the recording session, with no significant differences in whisking activity between control and DA-depleted mice (Q vs. W, p < 0.05 Supplementary 1A). During quiescent periods, no differences were observed in the resting membrane potential of dMSNs (control: -67.18 ± 1.30 mV; 6-OHDA: -67.89 ± 1.15 mV, p > 0.05, Figure 2D) although the input resistance increased (control: 120.10 ± 13.59 MΩ; 6-OHDA: 189.90 ± 1.15 MΩ, p < 0.05, Figure 2E). Interestingly, DA depletion significantly reduced spontaneous firing of dMSNs (control: 0.13 ± 0.07 Hz from 6 out of 16 neurons; 6-OHDA: 0 out of 14, p < 0.05, Figure 2F). In contrast, none of the previously mentioned properties changed in iMSNs (p > 0.15, Figure 2D, 2E, and 2F). Overall, our results show predominant changes in dMSNs membrane properties following DA depletion.

### MSNs depolarize before the onset of motor activity

We next measured the membrane potential dynamics during whisking in a subset of 48 MSNs, 23 in control mice (9 dMSNs and 14 iMSNs), and 25 in DA-depleted mice (8 dMSNs and 17 iMSNs) (Figure 3). In control mice, dMSNs, and iMSNs showed progressive depolarization that started approximately 50 ms before whisker movement onset, (Figure 3A). During whisking the membrane potential was more depolarized than during quiescent epochs in both MSN types (dMSN control Q: -68.10 ± 1.40 mV, W: -64.99 ± 1.38 mV, p < 0.01, Figure 3A, and 3C; iMSN control Q: -69.84 ± 1.32 mV, W: -67.62 ± 1.67 mV, p < 0.01, Figure 3A, and 3C). The peak depolarization was observed at 99 ± 14 ms in dMSNs, and at 101 ± 8 ms, in iMSNs after whisking onset. The membrane potential during whisking was as variable as during quiescent epochs, represented by the standard deviation (SD), in both dMSNs (dMSN control Q: 2.32 ± 0.41 mV, W: 1.70 ± 0.21 mV, p > 0.5, Figure 3A, and 3D) and iMSNs (iMSN control Q: 1.65 ± 0.17 mV, W: 1.47 ± 0.18 mV, p > 0.5, Figure 3A, and 3D). In DA-depleted mice, the whisking-induced depolarization was abolished in dMSNs but not in iMSNs (dMSN 6-OHDA Q: -66.88 ± 2.03 mV, W: -66.22 ± 2.00 mV, p > 0.05; iMSN 6-OHDA Q: -66.68 ± 1.12 mV, iMSN 6-OHDA W: -63.97 ± 1.21 mV, p < 0.01, Figure 3B-D). iMSNs in DA-depleted mice depolarized before whisking onset, as in control mice. The maximum peak of depolarization was observed 89 ± 15 ms after whisking onset. In DA-depleted mice, the membrane potential of dMSNs showed similar levels of variability during whisking and quiescence epochs, (dMSN 6-OHDA Q: 1.53 ± 0.14 mV, W: 1.20 ± 0.23 mV, p > 0.05, Figure 3B, and 3D) while in iMSNs, the membrane potential was less variable during whisking than in quiescence (iMSN 6-OHDA Q: 2.02 ± 0.25 mV, W: 1.59 ± 0.17 mV, p < 0.05, Figure 3B, and 3D). Our results show that MSNs in dorsolateral striatum encode and even predict whisker-related motor activity, and that DA depletion alters this coding in a pathway-dependent manner, reducing motor signals selectively in dMSNs.

**Figure 3.**
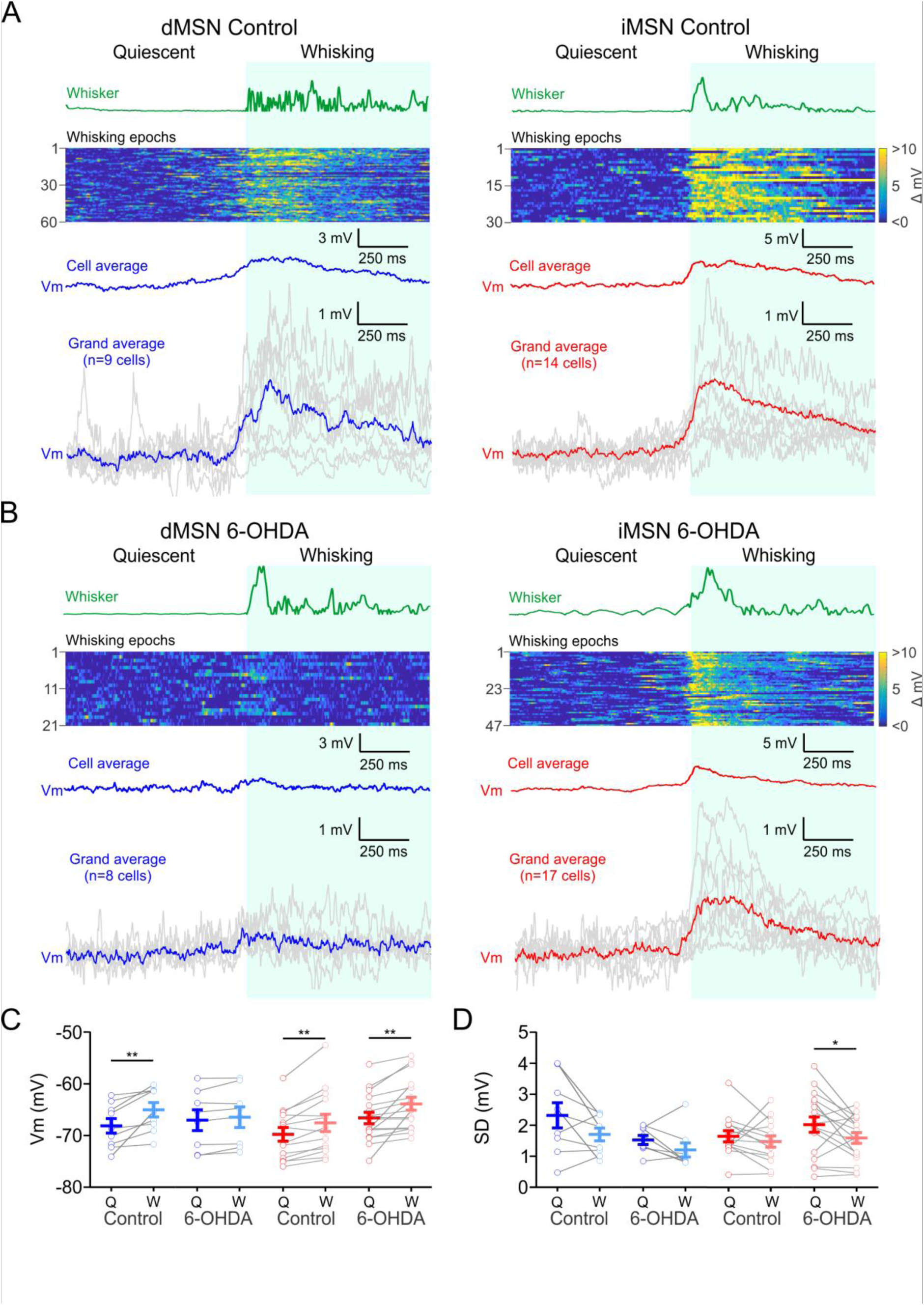
Whisker activity is preceded by membrane depolarization of MSNs in control conditions but not in dMSNs in DA-depleted striatum. ((A) Membrane potential depolarization of dMSN in control conditions precede whisking. Left from top to bottom: Example of whisker activity transition from quiescence to whisking (whisker, green). Heat map showing corresponding membrane potential fluctuations during repetitive whisking epochs in a single dMSN in control conditions (60 repetitions). The average membrane potential fluctuation for that cell is shown below (cell average, blue, scale bar as in the lesion example). Note that the depolarization starts a few milliseconds before the whisking movement. Grand average of the dMSNs; each grey trace indicates independent dMSN average activity (n = 9 cells). Right: Same as in the left panel but for iMSNs in control mice. Heat map (30 repetitions) and grand average (n=14 cells). (B) Same as in (A) but for 6-OHDA lesion mice. Heat maps (21 repetitions in dMSN and 47 in iMSN). Grand average (n=8 cells in dMSNs and n=17 cells in iMSNs). Note the absence of the depolarization during whisking in the dMSNs 6-OHDA in contrast with dMSNs in control conditions. (C) Membrane potential of dMSNs (blue) and iMSNs (red) during quiescent (Q) and whisking (W) epochs in control and 6-OHDA lesion mice. (D) Standard deviation (SD) of the membrane potential of dMSNs and iMSNs during Q and W epochs. For (C) and (D) each grey line represents the data from a single cell during Q (dark color) and W (light color), and error bars indicate mean ± SEM (blue). Paired t-test ^∗^ p < 0.05, ^∗∗^ p < 0.01.

### Sensory responses in MSNs are attenuated by whisking

Sensory-evoked responses represent a major source of synaptic input to MSNs in the DLS ^16,23^. In different cortical regions, both enhancement ^7,8,49^ and attenuation of sensory responses by motor activity was observed ^1,4,6^. Little is known about such sensorimotor interactions in the striatum. To examine whether and how whisking modulates sensory integration in MSNs, sensory stimuli were delivered as brief (15 ms) contralateral whisker deflections at random intervals (2 to 5 s). In parallel, whisker activity was monitored (Figure 4A, green line), allowing for the discrimination between stimuli delivered during whisker movement or quiescence (Figure 4A). Whisker deflections during quiescence evoked larger responses than those evoked during whisking in both dMSNs (dMSN control Q: 10.26 ± 3.09 mV; W: 5.31 ± 2.63 mV, p < 0.01, Figure 4B and 4D) and iMSNs (iMSN control Q: 7.94 ± 1.71 mV; W: 4.35 ± 0.74 mV, p < 0.05, Figure 4B and 4D). In contrast, no differences were observed in the latency of response peaks, neither in dMSNs (dMSN control Q: 33 ± 2 ms; W: 34 ± 4 ms, p > 0.05, Figure 4B and 4E) nor iMSNs (iMSN control Q: 31 ± 1 ms; W: 29 ± 2 ms, p > 0.05, Figure 4B and 4E). Differences between the amplitude of sensory responses during quiescence and whisking were maintained also in DA-depleted mice, in both dMSNs (dMSN 6-OHDA Q: 7.54 ± 1.42 mV; W: 4.62 ± 1.10 mV, p < 0.01, Figure 4C and 4D) and iMSNs (iMSN 6-OHDA Q: 8.90 ± 1.77 mV; W: 5.05 ± 1.47 mV, p < 0.001, Figure 4C and 4D). The temporal parameters were equally not affected following DA depletion, reaching peak depolarizations at similar times (Figure 4E, p > 0.05). In a subset of cells, ipsilateral responses were measured, also showing larger responses during quiescence than during whisking in both dMSNs and iMSNs, without difference in the temporal properties (Figure S2). One potential underlying mechanism for the observed response attenuation is an increase in membrane conductance of MSNs during whisking. Indeed, the input resistance of MSNs was lower during whisking, as measured by injection of current steps during quiescent and whisking periods (Figure S3). Our results show that whisking attenuates sensory responses in dMSNs and iMSNs, both under control conditions and in the DA-depleted striatum.

**Figure 4.**
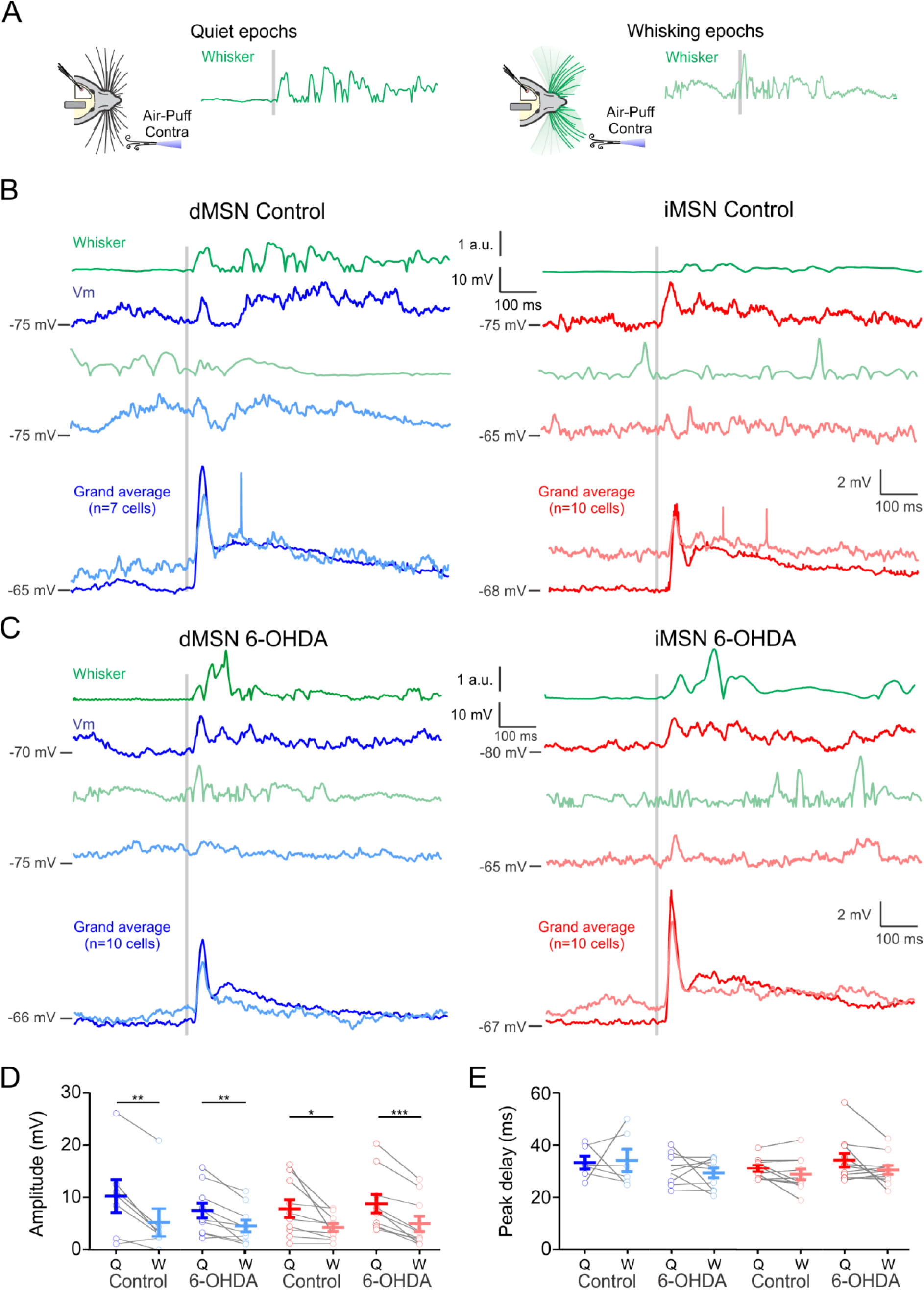
Sensory responses in MSNs are attenuated during whisking. ((A) Schematic of contralateral whisker stimulation during quiet epochs (left) or during whisking epochs (right). The stimulations were delivered randomly in intervals from 3 to 6 secs. Classification of events as occurring during Q or W was done post-hoc. The grey line indicates the trigger to stimulation. (B) Examples of traces from control dMSNs (blue left) and control iMSNs (red right) mice upon contralateral whisker stimulation (grey line). Dark color traces indicate the averaged response of individual MSNs to sensory stimulation during quiescence while light color traces indicate responses to sensory stimulation during whisking in the same cell. Whisker-related motor activity is indicated in green. Bottom panel, grand average of traces from independent MSNs in response to contralateral whisker stimulation in control mice (n=7 cells in dMSNs and n=10 cells in iMSNs). Note the decrease in amplitude in the sensory responses evoked during whisker movement in both MSN types. (C) Same as in (B) but for 6-OHDA lesion mice. Dark color traces indicate MSNs responses to sensory stimulation during quiescence while light color indicate responses to sensory stimulation during whisking. Grand average (n=10 cells in dMSNs and n=10 cells in iMSNs). (D) Contralateral whisker deflections produce larger amplitude responses in both MSN types during quiescence in control and 6-OHDA lesion mice (E) without affecting the peak of maximum depolarization (peak delay). For (D) and (E) each grey line represents the data from a single cell during Q (dark color) and W (light color), and error bars indicate mean ± SEM. Paired t-test ^∗^ p < 0.05, ^∗∗^ p < 0.01.

### DA depletion impairs bilateral discrimination in MSNs

We next examined how bilateral tactile stimuli are processed in DLS neurons, and whether bilateral responses are altered following DA depletion. To that end, we performed whole-cell recordings in head restrained awake mice while brief air puffs were delivered to the ipsi- or contralateral whiskers (Figure 5A). Ipsi- and contralateral whisker deflections delivered during quiescence evoked sensory responses in both dMSNs and iMSNs (Figure 5B and 5D). In control mice, contralateral whisker stimulation evoked a larger short-latency depolarization (within 50 ms from the stimulus onset) than ipsilateral stimulation, in both dMSNs and iMSNs (p < 0.05, Figure 5C, 5E, and Table S1). The onset delay of contralateral responses was also shorter than that of ipsilateral responses (p < 0.05, Figure 5C, 5E, and Table S1), although no differences were seen in the time of the peak amplitude (p > 0.05, Figure 5C, 5E, and Table S1). Following DA depletion, the differences between ipsi- and contralateral responses, both in terms of amplitude and onset latency, were abolished (p > 0.05, Figure 5C, 5E, and Table S1). These results show that DA depletion alters striatal sensory responses resulting in the diminished lateral encoding of tactile stimuli by MSNs.

**Figure 5.**
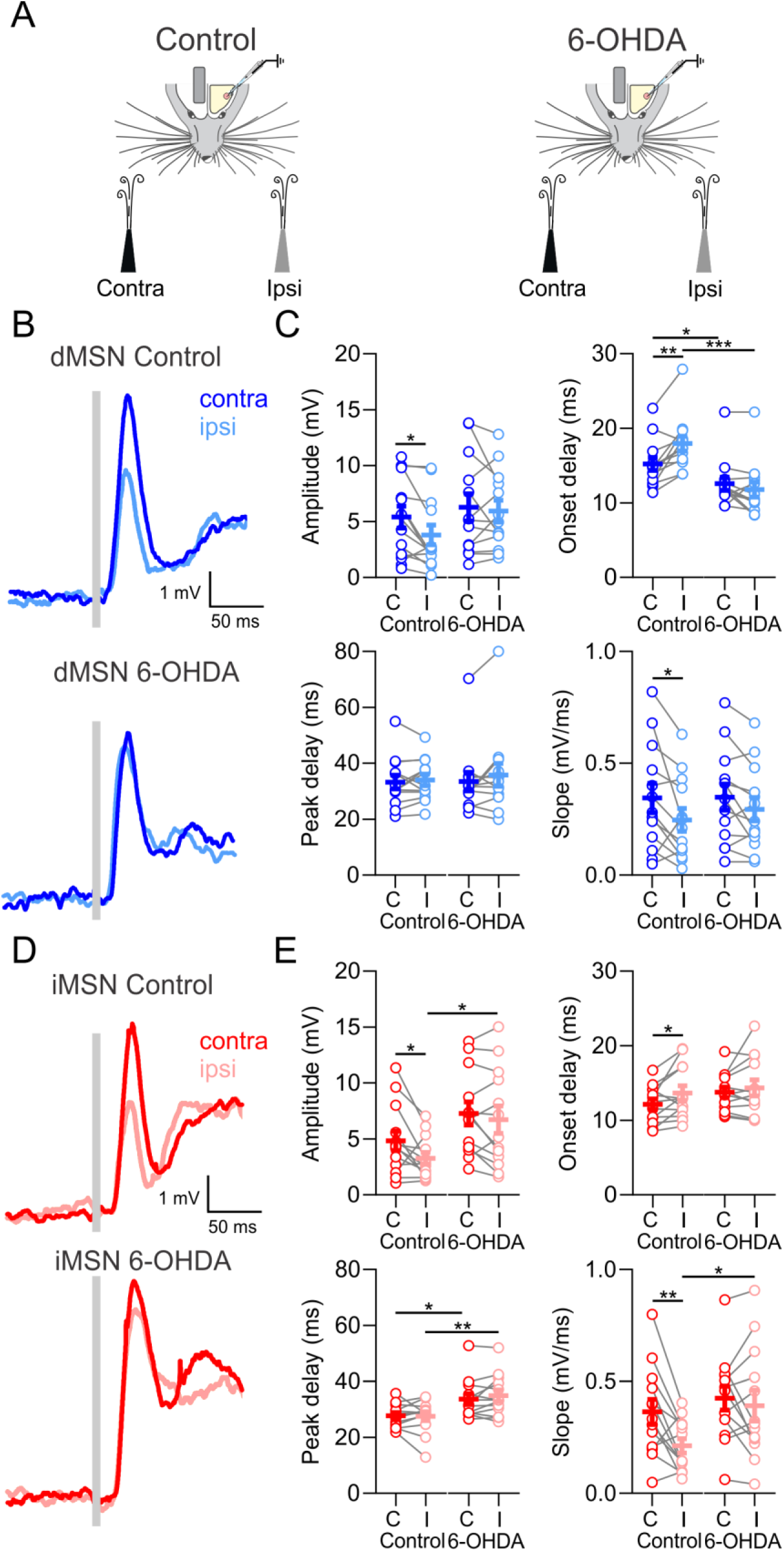
Laterality is encoded in the early component of sensory responses and impaired following DA depletion. ((A) Schematic of contralateral and ipsilateral whisker deflections during quiet epochs in control and 6-OHDA lesion mice. The stimulations were delivered randomly in intervals from 3 to 6 secs. (B) Grand average of responses of dMSNs in control (left) and 6-OHDA (right) lesion mice to contralateral (dark blue traces) and ipsilateral (light blue traces) whisker stimulation. The grey bar indicates the moment the stimulation was delivered. (C) Differences in response amplitude, onset delay, and slope of dMSNs between contralateral (C), and ipsilateral (I) responses (n = 13) are abolished in 6-OHDA lesion mice (n = 13). (D) Grand average of responses of iMSNs in control (left) and 6-OHDA (right) lesion mice to contralateral (red) and ipsilateral (light red) whisker stimulation. (E) Differences in response amplitude, onset delay, and slope of iMSNs between contralateral (C), and ipsilateral (I) responses (n = 13) are abolished in 6-OHDA lesion mice (n = 13). For (C) and (E) each grey line represents the data from a single cell during and error bars indicate mean ± SEM. Paired t-test * p < 0.05, ** p < 0.01, *** p < 0.001.

### DA depletion reduces the late sensory component in dMSNs

Sensory responses often had a secondary late component, between 100 and 250 ms after whisker deflection. Interestingly, this late component was more pronounced when stimuli were delivered during whisker quiescence, while it was almost absent when stimuli were delivered during whisking (Figure S2). The physiological relevance of this late component is not well understood. We tested whether the late component encodes stimulus laterality as the short-latency component (Figure 5), and if it is affected by DA depletion. In control mice, ipsi- and contralateral whisker stimulation produced late responses with no differences in the amplitude, onset, or area under the curve (AUC), in both dMSNs and iMSNs (p > 0.05, Figure 6C, 6E, Table S2). As in control mice, following DA depletion, no differences between ipsi- and contralateral responses were observed either in dMSNs or iMSNs (p > 0.05, Figure 6C, 6E, Table S2). However, the amplitude of the late response in dMSNs was significantly smaller in DA-depleted mice (p < 0.05, Figure 6C, and Table S2), which was not the case in iMSNs (p > 0.05, Figure 6E, and Table S2). These results indicate that although not encoding for the laterality of whisker deflections, the late component is affected in a cell-type-specific manner after DA depletion. One possible explanation for these differences could be the involvement of the late component in motor activity. Whisker deflection during quiescence often evoked whisking (QW trials), however, in a few cases the whiskers remained stationary after stimulation (QQ trials, Figure S5A). We compared the late component in QQ vs. QW trials in control and DA-depleted mice. Whisker stimulation in control mice evoked larger late responses in QW than in QQ trials in both MSN types (Figure S5). Following DA depletion, iMSNs still had larger responses during QW than those recorded during QQ. In contrast, the late component in dMSNs was reduced, and no differences were observed between QQ and QW trials. Thus, the late component is correlated with stimulus-evoked whisking in both MSNs types, but was reduced only in dMSNs following DA depletion. Our results show that in the DLS, the same individual MSNs represent both sensory inputs and motor activity. Sensory responses are attenuated by whisking, and dopamine plays a role in the representation of both processes.

**Figure 6.**
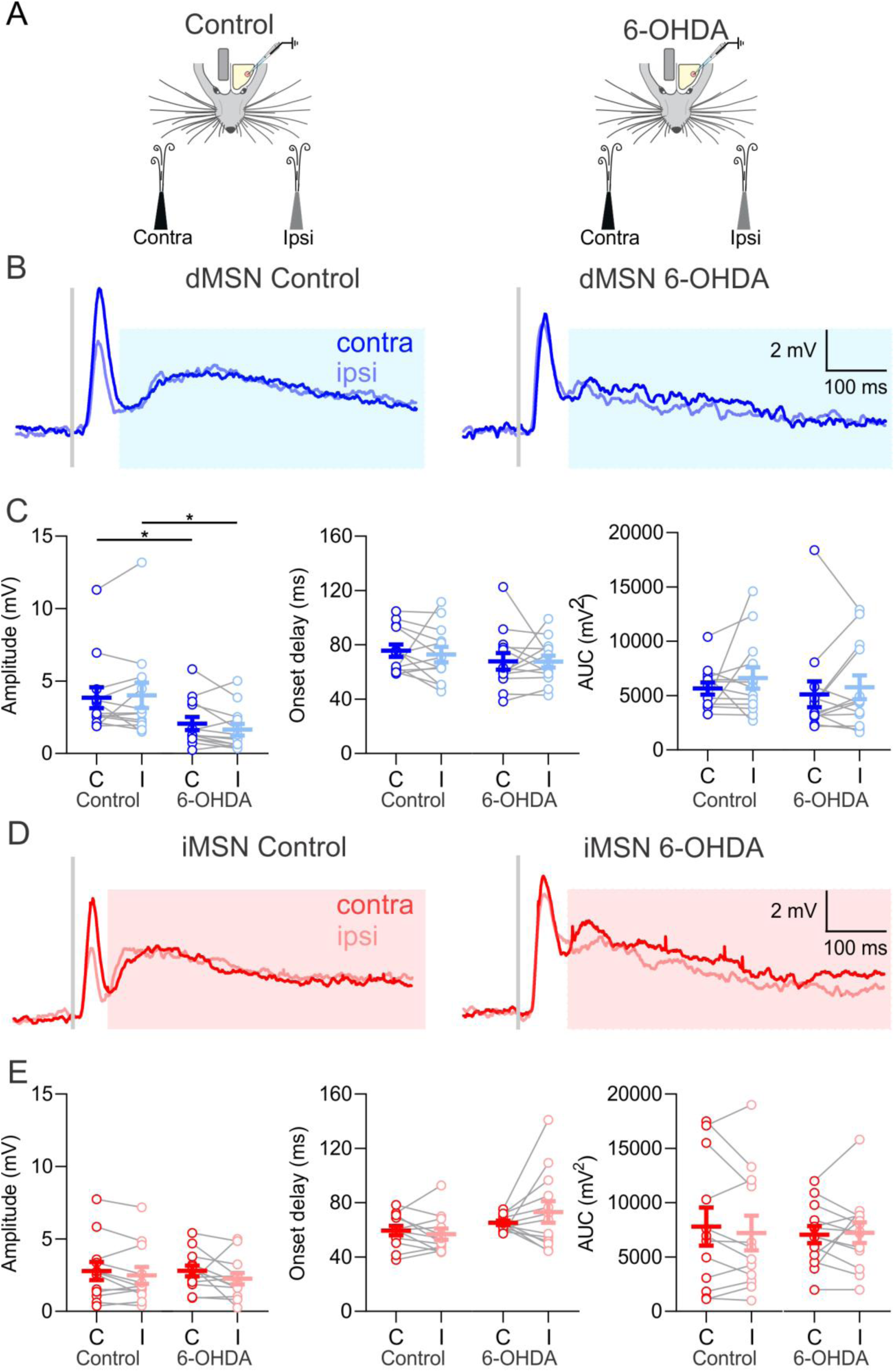
The late component of the sensory response is not involved in laterality encoding, and is reduced in dMSNs following DA depletion. ((A) Schematic of contralateral and ipsilateral whisker deflections during quiet epochs in control and 6-OHDA lesion mice. The stimulations were delivered randomly in intervals from 3 to 6 secs. (B) Grand average of the late component of the sensory response of dMSNs in control (left) and 6-OHDA (right) lesion mice to contralateral (dark blue traces) and ipsilateral (light blue traces) whisker stimulation. The light blue rectangle indicates the late component of the response to sensory stimulation. The grey bar indicates the moment the stimulation was delivered. Top middle: Schematic of contralateral and ipsilateral whisker deflections during quiet epochs in control and 6-OHDA lesion mice. The stimulations were delivered randomly in intervals from 3 to 6 secs. (C) No differences in response amplitude, onset delay or area under the curve (AUC) of dMSNs responses to contralateral (C) and ipsilateral (I) stimulations in control or 6-OHDA lesion mice. Note the diminution in response amplitude without affecting onset delay or area under the curve (AUC) (n = 13) in the dMSNs of 6-OHDA lesion mice compared to dMSNs in control mice (n = 13). (D) Grand average of the late component of iMSNs in control (left) and 6-OHDA (right) lesion mice to contralateral (red) and ipsilateral (light red) whisker stimulation. The light red rectangle indicates the late component of the response to sensory stimulation. The grey bar indicates the moment the stimulation was delivered. (E) No differences in response amplitude, onset delay or AUC of iMSNs between contralateral (C) and ipsilateral (I) stimulations in control (n=12) or 6-OHDA lesion mice (n=13). For (C) and (E) each grey line represents the data from a single cell during and error bars indicate mean ± SEM. Paired t-test ^∗^ p < 0.05, ^∗∗^ p < 0.01.

## Discussion

In this study we explored the modulation of striatal sensory processing by behavior and the changes resulting from DA depletion. Our data show that motor activity in both DA-intact and DA-depleted mice attenuated the responses to tactile stimulation in both striatal MSN types, and encoding of bilateral stimuli was impaired following DA depletion. Additionally, MSNs encoded spontaneous and evoked whisker movement and that DA depletion compromised both sensory and motor representations in a cell type-dependent manner.

### DA depletion affected motor integration preferentially in dMSNs

We showed that whisker movement was correlated with membrane potential depolarization in both dMSNs and iMSNs. Interestingly, depolarization preceded whisker movement onset, suggesting that the striatum is involved in preparatory neuronal activity occurring before movement initiation and execution. In agreement with previous studies that have shown co-activation of dMSNs and iMSNs ^50–54^, our data suggest a cooperative rather than antagonistic role during action initiation. The correlation between dMSN membrane potential and whisker movement was abolished following DA depletion. This effect was cell type-dependent and not observed in iMSNs. The attenuation of the late component of sensory responses in dMSNs (Figure 6 and S5) also points towards a cell type-specific effect of DA depletion. Together with the reduction in the spontaneous activity of dMSNs, our results show pathway-dependent changes in the striatal microcircuitry in PD, with reduction in the spontaneous and motor-related activity of the direct pathway. Supporting these results is a recent study using calcium imaging showing that a decrease in DA levels reduces the number of dMSNs recruited during locomotion ^54^.

### Sensory responses in both MSN types were attenuated by whisking

We show that both MSN types respond to bilateral whisker stimulation in awake mice. These results are in agreement with whole-cell recordings in anesthetized mice ^16,21,23^. A previous study using whole-cell recordings in awake mice reported that sensory responses in dMSNs were markedly larger than in iMSNs ^20^, which was not the case in our data. This discrepancy could be explained by differences in the experimental protocols. In the paper by Sippy and colleagues ^20^, only one whisker was stimulated while we delivered the sensory stimuli as an air-puff to several whiskers. Moreover, the small responses in iMSNs compared to dMSNs may reflect the learned task in that study. Previous studies report positive modulation for visual sensory processing during activated cortical states ^2, 7–9,49,55^ and negative modulatory effect for auditory sensory processing ^4–6^. We show an attenuation of striatal responses to whisker deflection during whisking. This reduction in the amplitude of sensory responses during active periods can be explained by a decrease in excitatory synaptic transmission due to adaptation in presynaptic structures during whisking ^1,56–58^. Another mechanism could be an increase in MSN membrane conductance due to synaptic input during whisking, and lastly, a decreased synaptic driving force due to membrane depolarization in MSNs. For instance, during up-states in anesthetized mice, a brain state characterized by depolarization and elevated synaptic input, sensory stimulation generates smaller responses in the MSNs than those evoked in the down-state ^16,21^. Our data show that during whisking, MSNs undergo depolarization and increase in membrane conductance (Figures 3 and S3), both supporting the attenuation of synaptic responses.

### DA depletion abolished laterality encoding in the early component of the sensory response

The early component of sensory responses (within 50 ms from the stimulus onset) has been linked to the initial whisker deflection ^16,20,22,23^. It also encodes information associated with laterality of whisker stimulation in anesthetized mice ^23^. We observed that MSNs responded to contralateral tactile inputs with earlier and larger depolarizing responses than to ipsilateral inputs. Whereas lateral encoding by MSNs is still present in awake mice, sensory responses were smaller and earlier than in anesthetized mice. Similar results were observed in the visual cortex, when comparing responses between anesthetized and awake mice ^59^. Following DA depletion, the bilateral asymmetry in the responses was abolished in both MSN types, rendering contralateral and ipsilateral sensory responses almost identical in terms of amplitude and latency. Cortical activity recorded as local field potentials in barrel cortex showed no changes in bilateral response asymmetry after DA depletion (Figure S4), suggesting that the loss of lateral encoding in MSNs is not an inherited effect due to upstream cortical changes. These results agree with previous results obtained from anesthetized mice^23^. What mechanisms could explain such effects of DA depletion? One possibility is that they are caused by differential changes in corticostriatal synapses originating from the pyramidal and intratelencephalic tracts. It was shown that DA depletion causes attenuation of MSNs responses to ipsilateral stimulation of the motor cortex ^36,60^. Another possible explanation for the lateral effects of DA depletion is the alteration of ipsilateral thalamostriatal inputs, ^61,62^.

### The late component of sensory responses did not encode stimulus laterality

The late component of the sensory response has been previously identified in both anesthetized and awake mice ^16,20,23^. Our data showed that ipsi- and contralateral whisker stimulation produced similar late responses in terms of amplitude and latency, suggesting no encoding of stimulus laterality. Moreover, we show that the late component was larger in cases where whisker deflection triggered whisking (Figure S5). Following DA depletion, the amplitude of the late component was reduced in dMSNs but not in iMSNs. In addition, dMSNs in DA-depleted mice showed no difference between trials that triggered whisking and those that did not. These results, and the observed decrease in whisking representation in dMSNs after DA depletion (Figure 3B), together suggest that the late component encodes sensory-evoked movement rather than sensory information per se.

In conclusion, we show that DLS neurons are involved in both sensory and motor processes, and that sensory processing is modulated by motor activity at the cellular level in both dMSNs and iMSNs. We also show that both sensory and motor representations are altered in the DA-depleted striatum, with motor processes being more affected in the direct pathway.

## METHODS

### Contact for Reagent and Resources Sharing

Further information and requests for resources and reagents should be directed to and will be fulfilled by the Lead Contact, Gilad Silberberg.

### Experimental Model and Subject Details

All experiments were performed according to the guidelines of the Stockholm municipal committee for animal experiments under an ethical permit to G.S. (N12/15). D1-Cre (EY217 line) or D2-Cre (ER44 line, GENSAT) or Adora2a-Cre (KG139, GENSAT) mice were crossed with the Channelrhodopsin (ChR2)-YFP reporter mouse lines (Ai32, the Jackson Laboratory) to induce expression of ChR2 in either dMSN or iMSN neurons. Mice of both sexes were housed under a 12 h light-dark cycle with food and water *ad libitum*.

## 6-OHDA Lesion

Adult mice of both sexes between 2 and 3 months of age were anesthetized with isofluorane and mounted in a stereotaxic frame (Stoelting). Mice received one unilateral injection of 1 µL of 6-OHDA-HCl (3.75 µg/µL dissolved in 0.02% ascorbic acid) into the MFB, at the following coordinates (in mm): antero-posterior -1.2, medio-lateral +1.2, and dorso-ventral -4.8. After surgery, all mice were injected with Temgesic (0.1 mg/Kg, Reckitt Benckiser Healthcare) and allowed to recover for at least 2 weeks. Only 6-OHDA-injected mice that showed rotational behavior were used in our experiments ^63^.

### Head implants

Adult mice (2 to 3 months old) were anesthetized with isofluorane, and the head was fixed in a stereotaxic apparatus. Temgesic (0.1 mg/Kg) was administered before the surgery. The body temperature was maintained at 36.5/37°C by a heating pad. An ocular ointment (Viscotears 2mg/g, Alcon) was applied over the eyes to prevent ocular dehydration. Lidocaine was applied on the skin surface before the incision. The skin covering the regions of interest was removed, and the bone gently cleaned. Targeted regions for intracellular and LFP recordings were marked using stereotaxic coordinates on the surface of the skull. Somatosensory cortex (S1) craniotomy coordinates for LFP: antero-posterior -1.5 mm to bregma, +3.2 mm medio-lateral to midsagittal suture. Striatum craniotomy coordinates for intracellular recordings: antero-posterior 0 mm, medio-lateral +3 mm. Then, a thin layer of light-curing adhesive (Ivoclar Vivadent) was applied on the exposed skull. An aluminum metal head-post was fixed with dental cement Tetric Evo (Ivoclar Vivadent) to the right hemisphere. A wall of dental cement was built along the edge of the bone covering the left hemisphere. After the surgery, the animals were returned to their home cage.

### *In vivo* field and whole-cell recordings

Experiments were performed as described previously ^20,23^. Following a recovery period of at least three days after implantation, mice were habituated to being head-restrained over a period of 3-4 days. On the day of the experiments, mice were anesthetized with isoflurane (3% for induction then 1.5%-2%), and small craniotomies (300-500 μm in diameter) were drilled to access the targeted areas. The open craniotomies were covered with silicone sealant (Kwik-Cast, WPI), and the animals were returned to their home cages for recovery. After 2 to 4 hours of recovery, mice were head-fixed, and the silicone from the craniotomies was removed. A bipolar tungsten electrode with an impedance of 1-2 MΩ was inserted 1 mm deep from the surface at the Barrel Field craniotomy (1.5 mm A-P; 3.25 mm M-L). Signals were acquired using a Differential AC Amplifier model 1700 (A-M Systems, USA) and digitized at 20 KHz with CED and Spike2 (Cambridge Electronic Design) parallel to whole-cell recordings. Patch-clamp recordings were performed in the DLS (0 mm A-P; 3 mm M-L) from 2 to 2.5 mm below the pia, a region that receives projections from sensory and motor areas (McGeorge and Faull, 1989). Patch pipettes were pulled with a puller P-1000 (Sutter Instruments). Pipettes (7-10 MOhm, borosilicate, Sutter) contained (in mM): 130 K-gluconate, 5 KCl, 10 HEPES, 4 Mg-ATP, 0.3 GTP, 10 Na2-phosphocreatine, and 0.2%–0.3% biocytin (pH = 7.2-7.3, osmolarity 280-290 mOsm). Signals were amplified using MultiClamp 700B amplifier (Molecular Devices) and digitized at 20 KHz with a CED acquisition board and Spike2 software (Cambridge Electronic Design). All patch-clamp recordings were obtained in current-clamp mode without current injection. Only for characterization of intrinsic electrophysiological properties, current injections from −110 pA to +110 pA in steps of 20 pA for 3 s each were applied. Values for quiet and whisking states for each current injection were extracted separately. Membrane potential was not corrected for liquid junction potentials. The optopatcher was used for the online identification of recorded MSNs ^46^ (A-M systems, WA USA). Light steps of 500ms were delivered at maximum LED power (3 mW at the tip of the fiber, 470 nm, Mightex systems) through an optic fiber inserted into the patch-pipette while recording the responses in whole-cell configuration. Positive cells responded to light pulses by a step-like depolarization, often exhibiting AP discharge, while negative cells did not show any response to light pulses.

### Whisking detection via reflector sensor

During whole-cell recordings, whisking epochs were detected by IR reflective sensor (HOA1405, 574 Honeywell, NC, USA)^64^. It was positioned pointing toward the whisker pad at a distance of 5 mm. The signal was acquired at 20 KHz using Spike2 software. Whisking epochs were defined as amplitudes bigger than 1.5 times the signal’s STD. Threshold crossing epochs with inter-epoch-interval smaller than 200 ms were binned together and defined as whisking epochs. Whisking epochs shorter than 500 ms were not considered for the analysis.

### Change in spontaneous activity in the quiet to whisking transition

Epochs including the transition from quiescence to whisking activity were selected, and one second time window was used to calculate the parameters of spontaneous activity for each behavior. After removing action potentials using a median filter, at least 10 epochs were considered to calculate the mean membrane potential and its variance. Heat maps were created with Matlab. All recordings were aligned to the onset of the movement.

### Whisker stimulation

Air puffs were delivered by picospritzer positioned 3 centimeters from the mouse’s whiskers edge. Randomized 15 ms air puff stimulations (ipsi- and contralateral) were delivered at a range of 0.2 to 0.33 Hz at least 10 responses were acquired for each stimulation condition. The air pressure was set to 15 psi. For the analysis, sensory responses evoked by air-puff stimulation were first sorted according to ipsi- or contralateral origin. Subsequently, these two groups were sorted according to the mouse behavior and then averaged to calculate the responses properties (amplitude, onset delay, peak delay, and slope) for quiescence or whisking.

### Cell labelling and immunohistochemistry

The cells were loaded with biocytin (Sigma) during the recording (>5 min recordings). At the end of the recording session, mice were sacrificed with an overdose of sodium pentobarbital (200 mg/kg I.P.) and transcardially perfused with 4% PFA in 0.01M PBS. The brain was extracted and kept for an additional 2 hr treatment in PFA, after which it was transferred to 0.01M PBS. Twenty-four hours before sectioning, the brain was transferred to and kept in 12% sucrose solution in 0.01M PBS. Coronal cryo-sections of 20 μm were mounted on microscope gelatin-coated slides and incubated overnight with Cy3 conjugated streptavidin (1:1000, Jackson ImmunoResearch Laboratories) in staining solution (1% BSA, 0.1% NaDeoxycholate, and 0.3% triton in 0.01 PBS) at 4°C. After washing in PBS, slides were mounted, and images were acquired on a fluorescence microscope in order to locate the recorded cell. In YFP positive mice, Cy3 positive cells were examined for YFP signal for classification as D1 or D2 expressing MSNs.

### TH Immunofluorescence

For TH immunofluorescence, coronal cryo-sections (20 µM) from the striatum were collected, mounted, rinsed in PBS and incubated in blocking solution (5% normal donkey serum, 0.3% Triton X-100, 1% BSA in PBS) for 30 min. Sections were incubated in anti-TH rabbit monoclonal antibody (Millipore) diluted 1:1000 in 1% PBS-BSA at 4°C overnight. The sections were rinsed in PBS and incubated for 2h at room temperature in Cy3-conjugated goat anti-rabbit secondary antibody (Jackson Laboratories) diluted 1:500 in 1% BSA-PBS. After processing, sections were examined with Zeiss Axio Imager M1 microscope (Carl Zeiss) and images of the dorsal striatum were captured at X4 magnification. TH staining showed over 80% reduction in fluorescence compared to the unlesioned hemisphere.

### Quantification and Statistical Analysis

All data are represented as mean ± SEM. All data distribution was first checked for normality (Shapiro-Wilk test) and analyzed accordingly. Normally distributed data were tested by one-way ANOVA followed by post hoc Tukey’s test analysis for multiple comparisons, and the unpaired and paired two-sample Student’s t-test was used for two-group comparisons. Non-normally distributed data were analyzed by the Kruskal-Wallis test for multi-group comparisons, followed by Mann-Whitney for two-group comparison, Wilcoxon signed-rank test was used for paired samples. Spontaneous firing proportions were analyzed using Fisher’s exact test. Statistical analyses were done in Prism and Matlab. Final figures have been assembled with Corel DrawX8.

## Acknowledgements

We thank Elin Dahlberg and Kristoffer Tenebro Berglund for technical assistance, Zach Chia, Matthijs Dorst, Johanna Frost-Nylen, Yvonne Johansson, and Anna Tokarska for helpful discussions. We thank Ramon Reig, Abdel El Manira, and Sten Grillner for comments on earlier versions of the manuscript and fruitful discussions. We are thankful to Ole Kiehn for the Ai32 mice and to Gilberto Fisone for the D2-Cre mice. This work was supported by a Wallenberg Fellowship from the Knut & Alice Wallenberg Foundation, Swedish Brain Foundation (Hjärnfonden), and Swedish Medical Research Council (VR-M) to G.S, R.d.l.T.M was supported by a Karolinska Institute postdoctoral scholarship.

## Author contribution

R.d.l.T.M , M.K. and G.S. conceived and planned the experiments. M.K. performed the 6- OHDA lesions, R.d.l.T.M. performed the experiments and analyzed the data. R.d.l.T.M, M.K. and G.S. wrote the manuscript.

## Declaration of interests

The authors declare no competing interests.

---

**Supplementary 1.**
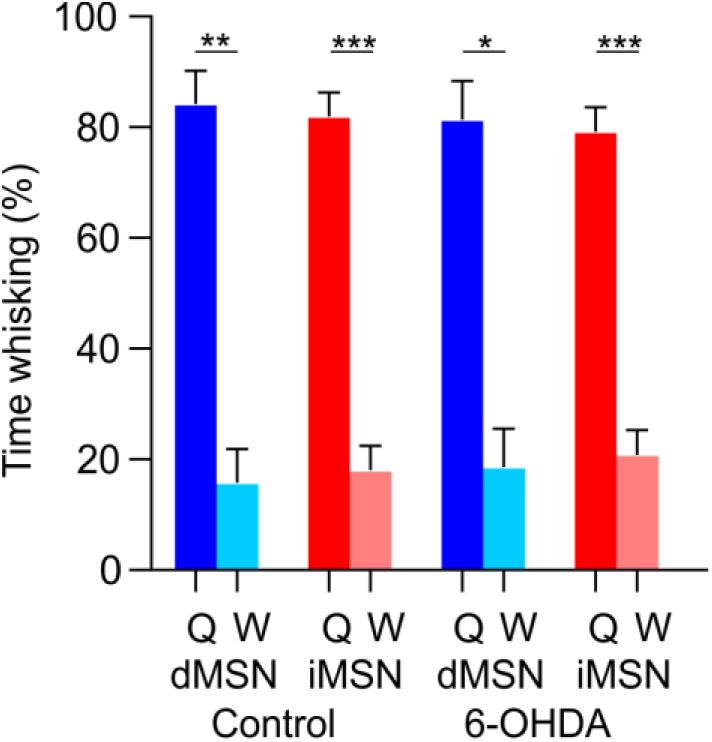
Fraction of time spent in quiescence and whisking during recordings performed in dMSNs and iMSN. (Data are given as the mean value ± SEM. Paired t-test and ANOVA. * p < 0.05, ** p < 0.01, *** p < 0.001.

**Supplementary 2 (Related to Figure 4).**
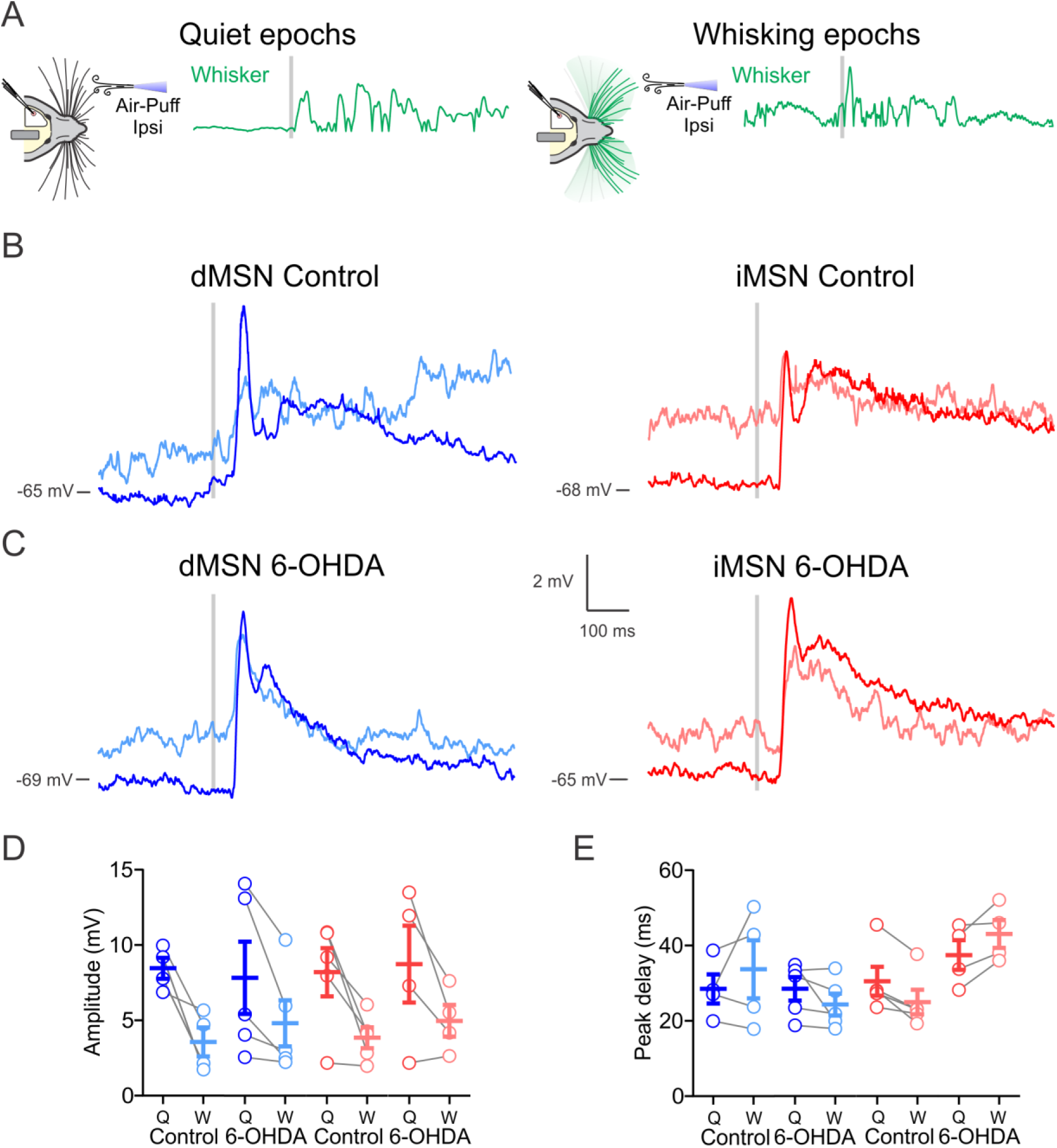
MSNs integrate ipsilateral tactile sensory information differently during quiescence and whisking in control and DA-depleted mice. ((A) Schematic of ipsilateral whisker stimulation during quiet epochs (left) or during whisking epochs (right). The stimulations were delivered randomly in intervals from 3 to 6 secs. Classification of events as occurring during Q or W was done post-hoc. The grey line indicates the trigger to stimulation. (B) Grand average of traces from control dMSNs (blue, left, n=4 cells) and control iMSNs (red, right, n=5 cells) mice upon ipsilateral whisker stimulation (grey line). Dark color traces indicate the averaged response of MSNs to sensory stimulation during quiescence, while light color traces indicate responses to sensory stimulation during whisking in the same cell. Note the decrease in amplitude in the sensory responses evoked during whisker movement in both MSN types. (C) Same as in (B) but for 6-OHDA lesion mice. Dark color trace indicates MSNs responses to sensory stimulation during quiescence, while light color indicates responses to sensory stimulation during whisking. Grand average (n=5 cells in dMSNs and n=4 cells in iMSNs). (D) Ipsilateral whisker deflections produce larger amplitude responses in both MSN types during quiescence in control and 6-OHDA lesion mice (E) without affecting the peak of maximum depolarization (peak delay). For (D) and (E), each grey line represents the data from a single cell during Q (dark color) and W (light color), and error bars indicate mean ± SEM.

**Supplementary 3. (Related to Figure 4).**
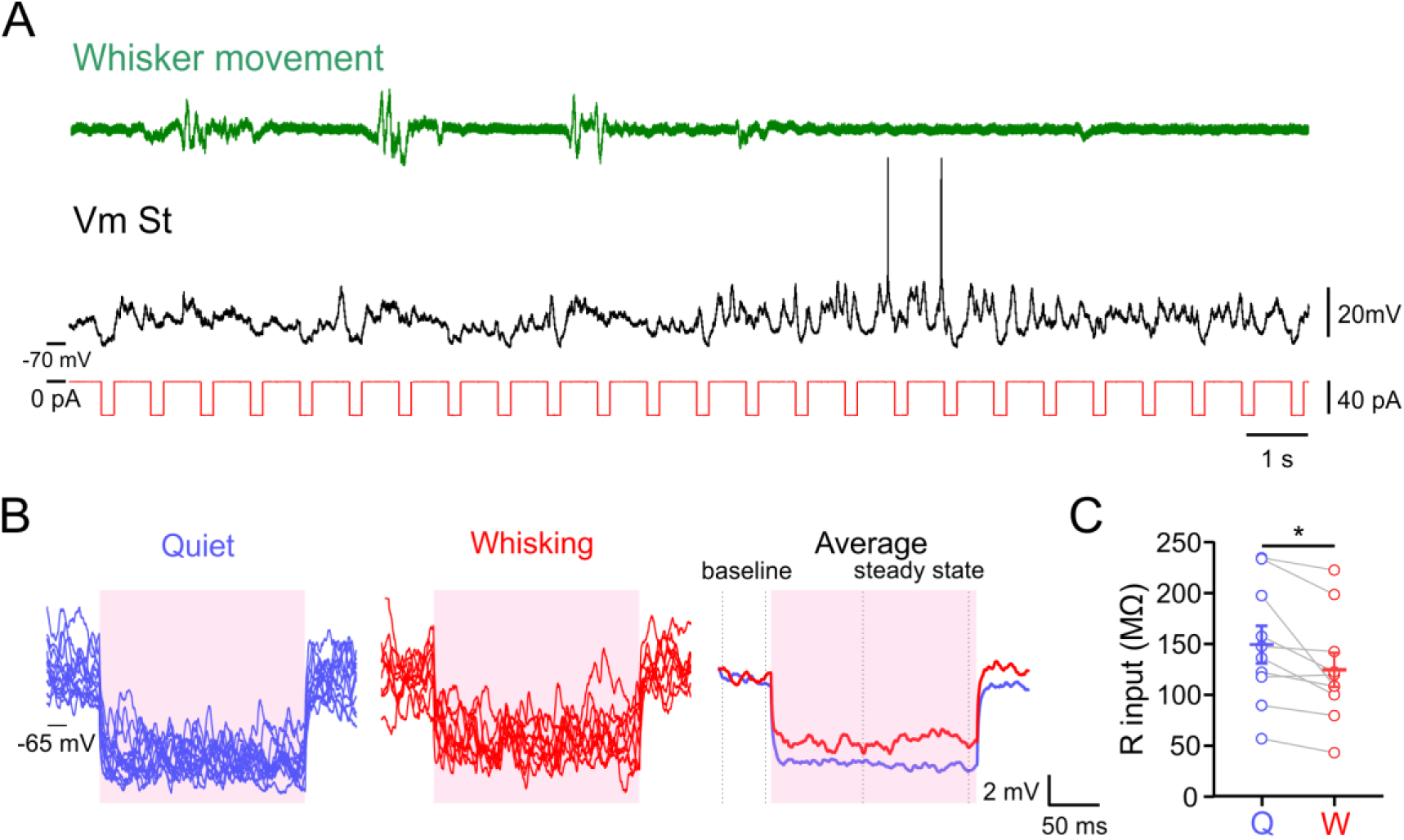
Input resistance of MSNs during quiescence and whisking in control mice. ((A) Example trace showing spontaneous membrane potential activity in an iMSN from DLS in control conditions of behaving mouse. Membrane potential (Vm, black) of the neuron was recorded simultaneously with the measurement of whisker activity (Whisker, green) while injecting square hyperpolarizing current pulses (-40 pA, 200 ms, red). (B) Current pulses occurring during quiescence (blue traces) and whisking (red traces) were averaged separately. The averages were vertically aligned to the same baseline. The R input was calculated using the difference between the average membrane potential 100 ms before the current pulse (baseline) and the average membrane potential for the last 100 ms of the current pulse (steady state). Trials that combine Q and W were discarded. (C) Input resistance during Q periods was significantly larger than during W epochs. Data are given as the mean value ± SEM. n=10. Paired t-test * p < 0.05.

**Supplementary 4.**
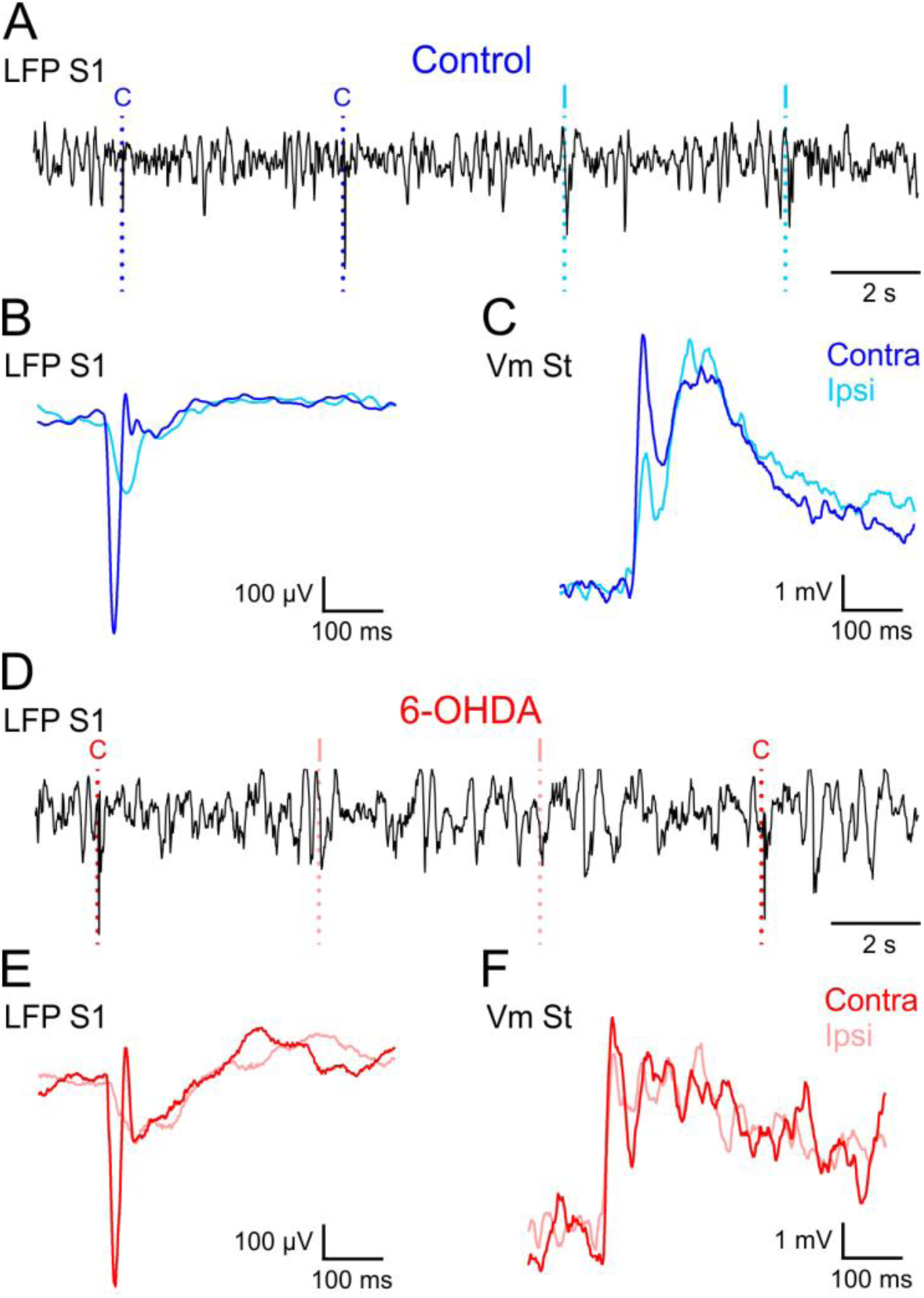
Laterality coding is impaired in the MSNs but not in the somatosensory cortex of DA-depleted mice. ((A) Example traces showing LFP activity in somatosensory cortex in control mice. A dashed line indicates the contralateral (dark blue, C) and ipsilateral (light blue, I) onset of the whisker stimulations. (B) Average of LFP responses of somatosensory cortex in control mice to contralateral (dark blue traces) and ipsilateral (light blue traces) whisker stimulation. (C) Average of responses of MSN in control mice to contralateral (dark blue traces) and ipsilateral (light blue traces) whisker stimulation. Both B and C were recorded in parallel and depict the average responses of cortex and striatum to the same whisker stimulations. (D) Example traces showing LFP activity in somatosensory cortex in 6-OHDA mice. A dashed line indicates the contralateral (dark red, C) and ipsilateral (light red, I) onset of the whisker stimulations (E) Average of LFP responses of somatosensory cortex in 6-OHDA mice to contralateral (dark red traces) and ipsilateral (light red traces) whisker stimulation. (F) Average of responses of MSNs in 6-OHDA mice to contralateral (dark red traces) and ipsilateral (light red traces) whisker stimulation. Both E and F were recorded in parallel and depict the average responses of cortex and striatum to the same whisker stimulations.

**Supplementary 5.**
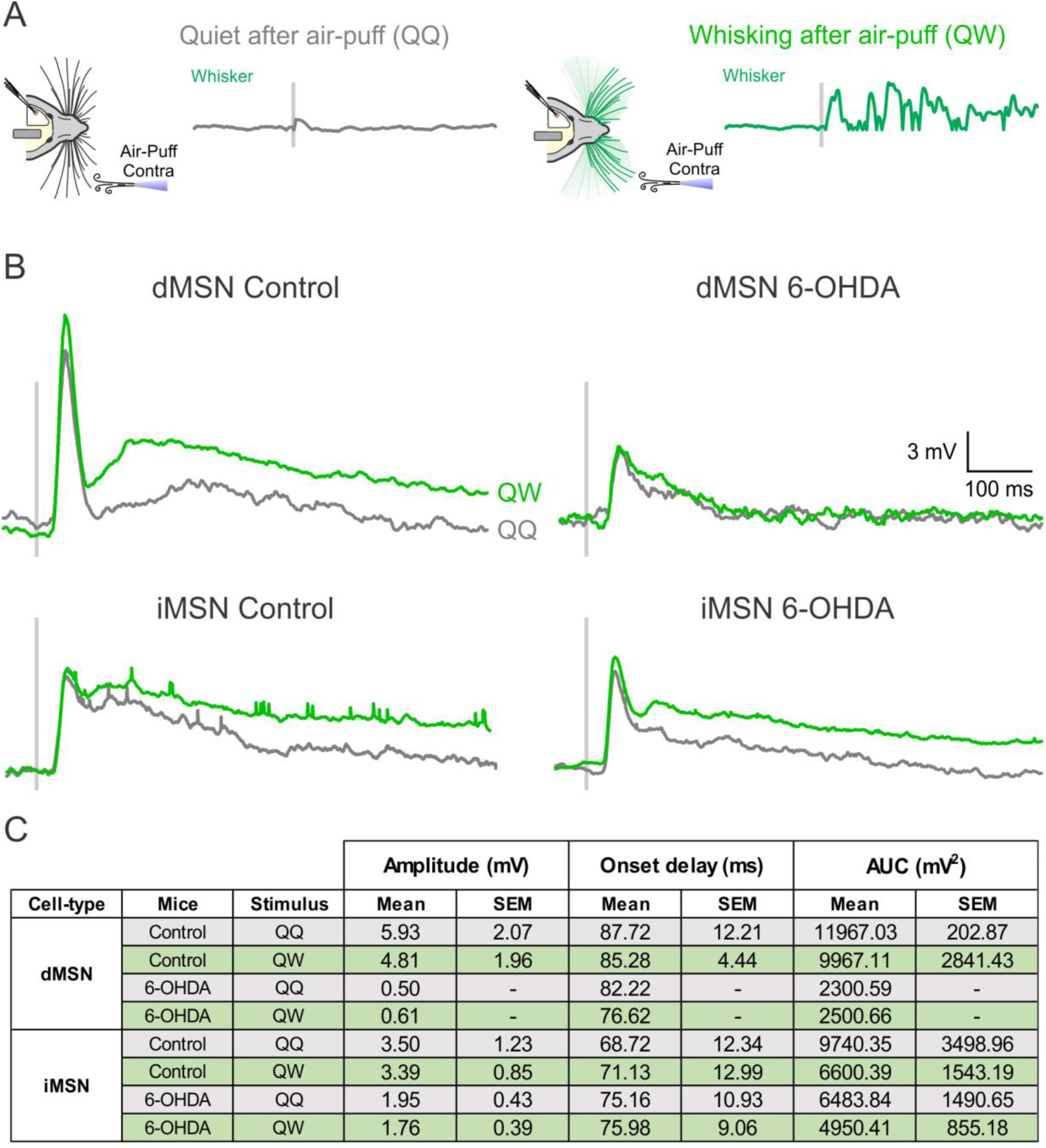
The late component of the sensory response correlates with whisker movement and is reduced in dMSNs following DA depletion. ((A) Schematic of contralateral whisker deflections during quiet epochs that do not promote whisker movement after the stimulation (QQ, left) and whisker deflections during quiet epochs that trigger whisker movement after the stimulation (QW, right). The grey line indicates the moment the stimulation was delivered. (B) Left: Grand average of membrane potential traces from dMSNs and IMSNs in control mice upon contralateral whisker deflections during QQ (grey trace) and whisker deflections during QW (green trace). Right: Same as in the left panel but for dMSNs and iMSNs in 6- OHDA lesion mice. control dMSN n= 3, control iMSN n= 5, 6-OHDA dMSN n= 1, 6-OHDA iMSN n= 6. (C) Late component of the tactile sensory response recorded in QQ and QW conditions in identified MSNs from control and 6-OHDA mice.

**Table S1 (related to Figure 5):** Short-latency responses to whisker stimulation in identified MSNs from control and DA-depleted mice.

| Cell-type | Mice | Stimulus | Amplitude (mV) |  | Onset delay (ms) |  | Peak delay (ms) |  | Slope (mV/ms) |  |
| --- | --- | --- | --- | --- | --- | --- | --- | --- | --- | --- |
|  |  |  | Mean | SEM | Mean | SEM | Mean | SEM | Mean | SEM |
| **dMSN** | control | Contra- | 5.55 | 0.98 | 15.20 | 0.87 | 33.22 | 2.46 | 0.34 | 0.07 |
|  | control | Ipsi- | 3.95 | 0.88 | 17.94 | 0.99 | 34.01 | 2.08 | 0.25 | 0.05 |
|  | 6-OHDA | Contra- | 6.27 | 1.22 | 12.54 | 0.89 | 33.45 | 3.31 | 0.35 | 0.06 |
|  | 6-OHDA | Ipsi- | 5.93 | 0.97 | 11.77 | 1.01 | 36.03 | 4.28 | 0.29 | 0.05 |
| **iMSN** | control | Contra- | 4.84 | 0.87 | 12.14 | 0.69 | 27.70 | 1.13 | 0.34 | 0.06 |
|  | control | Ipsi- | 3.27 | 0.52 | 13.64 | 0.98 | 27.55 | 1.59 | 0.23 | 0.03 |
|  | 6-OHDA | Contra- | 7.26 | 1.03 | 13.74 | 0.71 | 33.70 | 1.96 | 0.43 | 0.05 |
|  | 6-OHDA | Ipsi- | 6.71 | 1.22 | 14.32 | 1.09 | 34.90 | 2.06 | 0.39 | 0.07 |

**Table S2 (related to Figure 6):** Late responses to whisker stimulation in identified MSNs from control and DA-depleted mice.

| Cell-type | Mice | Stimulus | Amplitude (mV) |  | Onset delay (ms) |  | AUC (mV<sup>2</sup>) |  |
| --- | --- | --- | --- | --- | --- | --- | --- | --- |
|  |  |  | Mean | SEM | Mean | SEM | Mean | SEM |
| **dMSN** | control | Contra- | 3.85 | 0.73 | 75.77 | 4.57 | 5651.54 | 559.53 |
|  | control | Ipsi- | 4.01 | 0.87 | 72.87 | 5.65 | 6613.08 | 985.42 |
|  | 6-OHDA | Contra- | 2.06 | 0.45 | 67.83 | 6.15 | 5113.08 | 1190.29 |
|  | 6-OHDA | Ipsi- | 1.64 | 0.40 | 67.72 | 4.38 | 5767.69 | 1092.77 |
| **iMSN** | control | Contra- | 2.78 | 0.63 | 59.43 | 3.53 | 7803.33 | 1746.24 |
|  | control | Ipsi- | 2.48 | 0.58 | 56.88 | 4.17 | 7212.27 | 1589.20 |
|  | 6-OHDA | Contra- | 2.79 | 0.36 | 65.21 | 1.50 | 7062.31 | 785.27 |
|  | 6-OHDA | Ipsi- | 2.26 | 0.39 | 73.14 | 7.95 | 7242.31 | 956.92 |

## Notes

### Competing Interest Statement

The authors have declared no competing interest.

